# Refinement of a technique for collecting and evaluating the osmolality of haemolymph from *Drosophila* larvae

**DOI:** 10.1101/2023.12.25.573325

**Authors:** Misato Kurio, Yuma Tsukasa, Tadashi Uemura, Tadao Usui

**Author notes:** Correspondence (T.Us.).

## Abstract

*ex vivo* physiological experiments using small insect models such as *Drosophila* larvae have become increasingly useful to address fundamental biological questions. To perform such experiments, various artificial saline solutions have been developed, but their osmolality varies significantly from one to the next. Such a variation of osmolality stems, in part, from the difficulty of determining the true value of haemolymph osmolality in *Drosophila* larvae. Thus, there is a pressing need to refine protocols for collecting and measuring the osmolality of the larval haemolymph. Two major obstacles are thought to impede the accurate analysis of haemolymph collected from small insects: melanin formation and gut-derived contamination. Here, we greatly refined existing haemolymph collecting methods, evaluated the purity of the collected haemolymph under melanin-free conditions, and concluded that the true value of haemolymph osmolality is close to 306.0 mOsm kg^−1^ in *Drosophila* larvae.

## INTRODUCTION

In recent years, *ex vivo* physiological experiments using small insect models such as *Drosophila* larvae have become increasingly important (Handke et al., 2014; Hu et al., 2020; Kanaoka et al., 2023; Lemon et al., 2015; Li et al., 2023; Nakamizo-Dojo et al., 2023; Onodera et al., 2017; Redhai et al., 2020; Terada et al., 2016). To perform such experiments, various artificial saline solutions have been developed. Notably, the osmolality settings are significantly different between solutions: the normal saline solution (also known as Standard saline, 300.0 ∼ 315.5 mOsm kg^−1^, Jan LY & Jan YN., 1976); the external saline solution (305.0 mOsm kg^−1^, Xiang et al., 2010); Baines external solution (310∼320 mOsm kg^−1^, Jovanic et al., 2016); Haemolymph-Like No.3 (HL3, 343.0 mOsm kg^−1^; Stewart et al., 1994); HL3.1 (309.1 mOsm kg^−1^, Feng et al., 2004); HL4/HL5 (361.0 mOsm kg^−1^, Stewart et al., 1994); HL6 (341.0 mOsm kg^−1^, Macleod et al., 2002); and Schneider’s *Drosophila* medium (360∼374 mOsm kg^−1^, Schneider, 1972). These variable settings of osmolality rely on the single reported value of osmolality (390 ± 5.5 mOsm kg^−1^; Pierce et al., 1999). Therefore, once another reliable value of haemolymph osmolality is determined, the saline solution can be further optimized accordingly. Toward this objective, it is critically important to refine protocols for collecting pure haemolymph and measuring its osmolality.

In relatively large insects, such as tobacco hornworm larvae and adult honeybees, a substantial volume of fluid (≥ 0.2 mL) can quickly be collected without contamination, as it leaks spontaneously from an incised site on a leg or antenna (Adams & Wilcox, 1973; Borsuk et al., 2017; Łoś & Strachecka, 2018). The osmolality of haemolymph can be assessed by analyzing either freezing-point or vapor-pressure depression. In contrast, for small insects, the haemolymph volume in each animal is extremely limited, necessitating the simultaneous processing of dozens of animals to obtain enough for osmolality measurement (≥ 10 μL). Although a capillary-type osmometer requires only a tiny amount of liquid (≥ 2 nL) for measurement, it is still difficult to extract haemolymph from legless animals like *Drosophila* or honeybee larvae without contamination. In addition, two major obstacles hinder the accurate measurement of the haemolymph collected from small insects: (1) various biochemical reactions, such as melanization, are initiated as it takes longer to collect haemolymph from a batch of animals; (2) gut-derived contents contaminate the collected haemolymph (Borsuk et al., 2017; Palomino-Schätzlein et al., 2022; Tabunoki et al., 2019). Specifically, the melanization reaction yields heterogeneous biopolymers, such as eumelanin and pheomelanin, by consuming organic solutes in haemolymph like tyrosine and cysteine, thereby likely reducing its osmolality. On the other hand, when the haemolymph is contaminated by gut-derived materials, the measured osmolality should spuriously increase because soluble components of intestinal contents are highly concentrated in the hindgut lumen (Harpur & Popkin, 1965).

In our present study, we aimed to establish a more reliable and reproducible osmolality measurement protocol for small insects. To this end, we reexamined each step of the preceding haemolymph collecting methods (Palomino-Schätzlein et al., 2022; Pierce et al., 1999) and evaluated the purity of the collected haemolymph by employing a dye-tracing technique to detect food-derived materials (Sano et al., 2015), while haemolymph melanization was sufficiently suppressed by introducing a specific genetic mutation (Nam et al., 2012).

## MATERIALS AND METHODS

### Fly husbandry

The flies were raised on a standard cornmeal-yeast-agar food at 25°C in constant-dark conditions. The following stocks were used: *y*^1^ *w*^67c23^; +; *P{CaryP}attP2* (BDSC 8622) and *Hayan*^1^ (a gift from Won-Jae Lee, Seoul National University, Republic of Korea; Nam et al., 2012).

### Collection of haemolymph using centrifugation and measurement of its osmolality

For tracing food-derived materials, 10 g L^−1^ Brilliant Blue FCF (027-12842, Wako Pure Chemical Industry, Japan) was added to the food. For analysis, we crossed 20 female and 10 males in a food vial without blue dye for three days, and then transferred these flies to a new food vial with blue dye. In this vial, the flies laid eggs, and then the hatched larvae fed on the blue dye-labeled food.

Before starting each experiment, the osmometer was calibrated carefully. 60 third-instar wandering larvae were recovered from vials with a paint brush. Subsequent operations up to centrifugation were performed in temperature-controlled rooms, at 4 or 25°C (cold temperature [CT] and room temperature [RT], respectively), and all instruments and reagents were precooled to the respective temperatures before starting the procedures. Larvae were quickly washed twice in a 35-mm plastic dish (1000-035, AGC TECHNO GLASS Co., Ltd., Japan) filled with pure water (06442-95, NACALAI TESQUE, INC., Japan) and transferred onto a sheet of KimWipes® absorbent paper (S200 62011J-240, NIPPON PAPER CRECIA Co., Ltd., Japan), and excess water was removed with a dry brush. Larvae were processed in groups of 30 individuals at a time. Under a binocular dissecting microscope, the dorsal epidermis of the second thorax (T2) was carefully torn with tweezers (0208-5-PS, Dumont, Switzerland) to make a small slit, and 30 larvae were then carefully transferred to two separate 0.2-mL PCR tube (NN-2719S, TERUMO Corp., Japan) slightly off-center at the bottom. The 0.2-mL PCR tube was then placed in a 1.5-mL tube (131-615c, Fukae-kasei Co., Ltd, Japan) and centrifuged at 500 G for 1 minute at 4 or 25°C using a refrigerated microcentrifuge (MX-300, TOMY, Japan). Next, the first PCR tube was discarded, and the second batch of 30 larvae, processed similarly and placed in a punctured 0.2-mL PCR tube, was placed in the same 1.5-mL tube and centrifuged, thus combining the haemolymph from all 60 larvae. After centrifugation, the second PCR tube was discarded. Importantly, for experiments under CT condition, the second PCR tube was chilled at 4°C until just before centrifugation. The 0.2-mL PCR tube was set so that the hole in the bottom faced vertically downward when the tube was placed in a fixed angle rotor. After confirming that the osmometer was calibrated correctly using the 290 mOsm kg^−1^ standard solution (this procedure usually takes 2–3 minutes), an aliquot of 10 μL supernatant from the collected haemolymph in the 1.5-mL tube was transferred to the osmometer (VAPRO 5520, Wescor). It should be noted that the value of osmolality is expressed in “mmol kg^−1^” when using this osmometer, which is the same as the value expressed in “mOsm kg^−1^”.

### Evaluation of food-derived materials in haemolymph

The collected haemolymph was incubated overnight to complete the melanization reaction at RT. A 2 μL aliquot of haemolymph was then used for an optical absorbance spectrum from 190 to 840 nm with a Nanodrop 2000c spectrometer (ND-2000c, Thermo Fisher Scientific Inc., MA), employing pure water as a reference for absorbance measurements.

### Collection of haemolymph without centrifugation and evaluation of its food-derived materials

Ten incised larvae were prepared as described above, and were then piled up on a strip of PARAFILM® (P7543-1EA, Sigma-Aldrich, MA). The leaking haemolymph was transferred into a 1.5-mL tube using a micropipette. The absorbance was measured immediately after collecting the haemolymph.

### Data analyses

All data analyses were performed using Python 3 (version 3.7.6, Python Software Foundation) and R (version 3.6.1, R Development Core Team). Using Python packages, a linear model was applied to determine how the fluid osmolality correlated with contamination by gut-derived materials. To estimate the true value of haemolymph osmolality, we obtained the *Y*-intercept of the linear equation using LinearRegression() in the Python package sklearn.linear_model. A Spearman’s rank correlation test was performed to examine the correlation between absorbance at 629 nm and osmolality of haemolymph using cor.test(). 95% confidence intervals were obtained using Python packages. A Mann-Whitney *U*-test was performed using wilcox.exact() in the R package exactRanktests (version 0.8.31).

## RESULTS AND DISCUSSION

To minimize the melanization reaction that could potentially affect the osmolality of the collected haemolymph, we utilized a *Drosophila* strain with a loss-of-function *Hayan* mutation (*Hayan*^1^; Nam et al., 2012); *Hayan* encodes a serine protease that cleaves pro-phenoloxidase to yield phenoloxidase, a key melanin-producing enzyme that is active during microbial infection or after epidermal injury (Dudzic et al., 2019; Nam et al., 2012). In addition, we labeled the larval food with blue dye (1g L^−1^ Brilliant blue FCF, see details in MATERIALS AND METHODS) to trace the intestinal contents of food origin during haemolymph collection.

By tearing the dorsal epidermis of a *Drosophila* third instar wandering larva at room temperature (RT, 25.0 ± 2.0°C), we collected haemolymph leaking from the incised site (Figure 1), and found some contaminating bluish material. To examine the fluid osmolality and evaluate the potential contaminants, we then collected more than 20 μL of haemolymph by centrifugating the incised larvae (60 larvae; Palomino-Schätzlein et al., 2022; see details in MATERIALS AND METHODS). We measured the osmolality and the absorbance spectrum (190 ∼ 840 nm) of the collected fluid, in which the absorbance at 629 nm specifically indicates the presence of the blue dye (Figure 2 and 3). The optical absorbance at 629 nm was 1.6 ± 0.3 (AU), and the measured osmolality was 352 ± 6.6 mOsm kg^−1^, indicative of the contamination by food-derived materials (*n* = 7; Figure 2A). Notably, the moderate magnitude of the absorbance in the range from 300 to 600 nm possibly originated from the intrinsic optical properties of the food-derived materials and/or melanin pigments produced by residual phenoloxidase activity in the *Hayan*^1^ mutant (Dudzic et al., 2019). These results suggest that the measured osmolality could be markedly overestimated due to contamination by food-derived materials under RT conditions.

**Figure 1.**
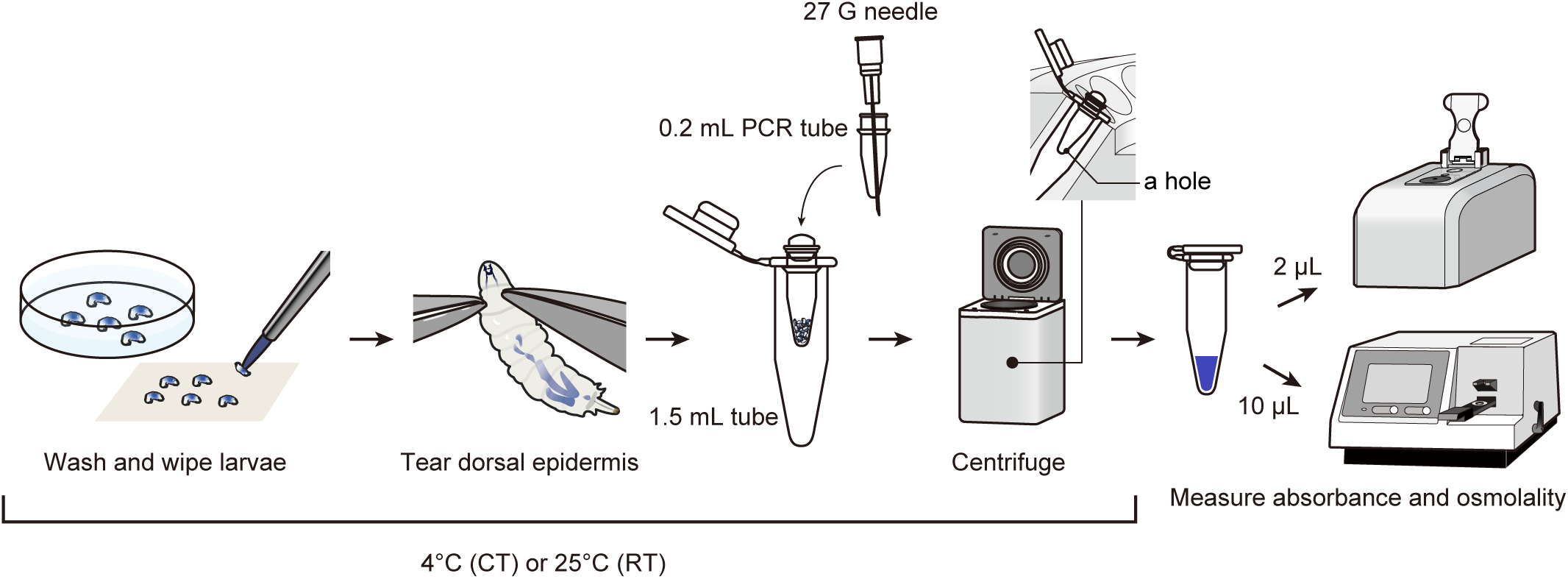
Overview of the refined method for collecting haemolymph from *Drosophila* larvae by centrifugation.

**Figure 2.**
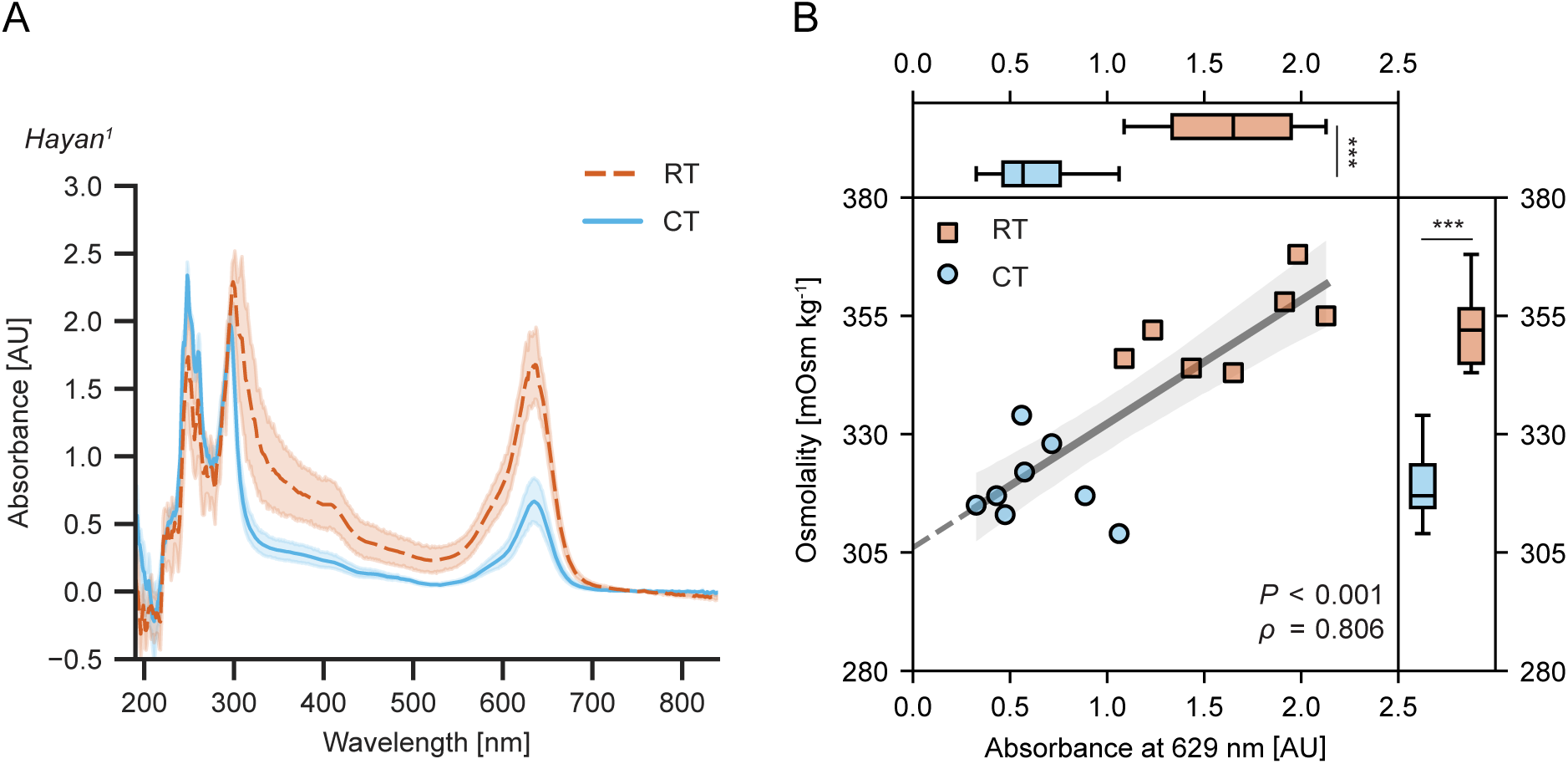
The purity and osmolality of haemolymph collected from *Hayan*^1^ mutant larvae by centrifugation depends on temperature conditions. (A) Absorption spectra of haemolymph collected from *Hayan*^1^ mutant larvae at 25°C (room temperature (RT) condition) or 4°C (cold temperature (CT) condition). The orange dashed line and solid blue line indicate mean values of absorbance under RT or CT condition, respectively. Orange and blue shading indicates the 95% confidence intervals of absorbance under RT or CT conditions, respectively. (B) Absorbance at 629 nm and osmolality of haemolymph collected from *Hayan*^1^ mutant larvae under RT or CT conditions are displayed in scatter plots and box plots. \**P* < 0.05, \*\**P* < 0.01, \*\*\**P* < 0.001, Mann-Whitney *U*-test; *n* = 7 in RT and *n* = 8 in CT. The dark gray line indicates linear regression. Light gray shading indicates the 95% confidence interval.

**Figure 3.**
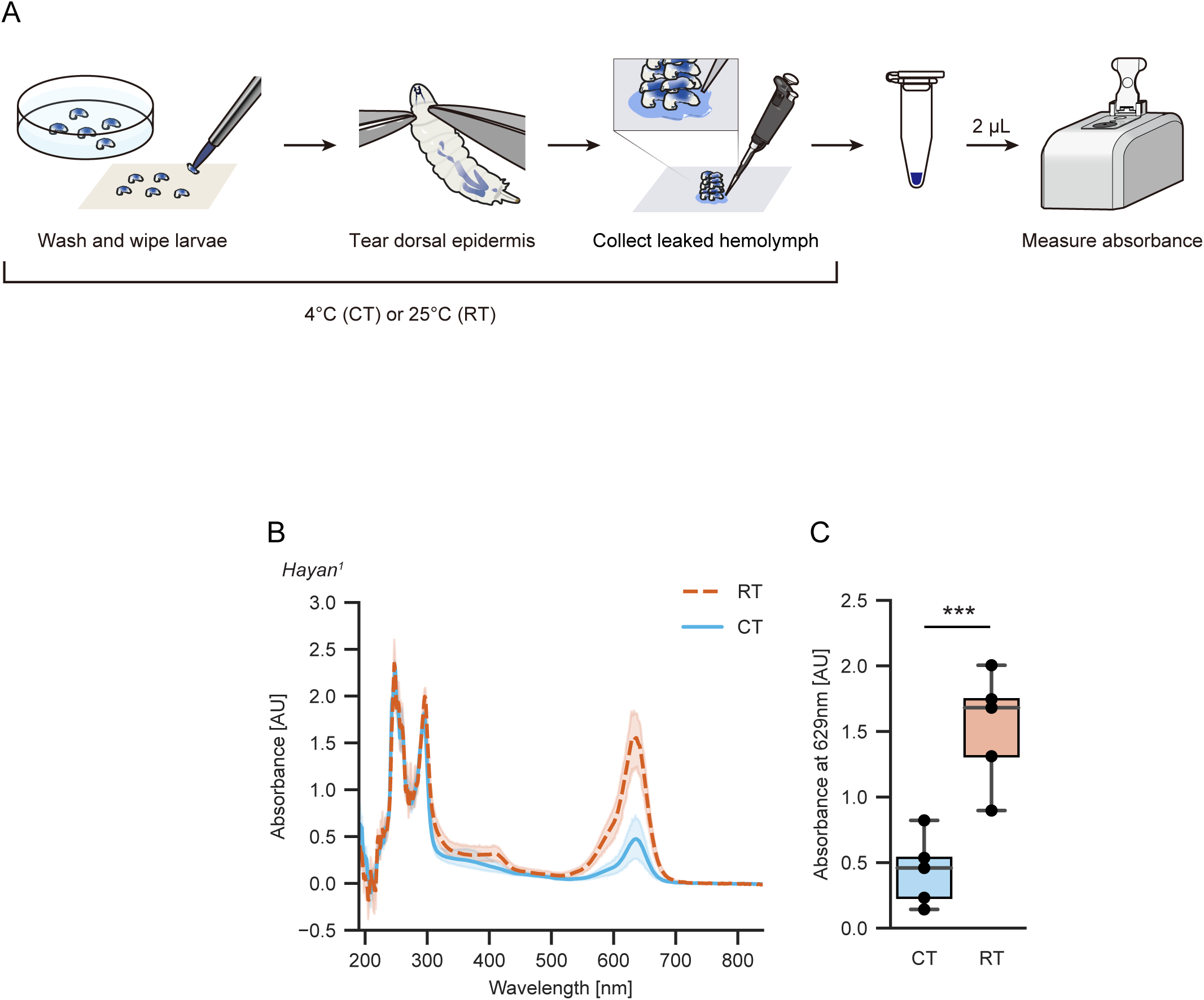
Gut-derived materials contaminate samples before the centrifugation step under RT conditions. (A) Overview of a method for collecting haemolymph leaking spontaneously from *Drosophila* larvae without centrifugation. (B) Absorption spectra of haemolymph collected without centrifugation from *Hayan*^1^ mutant larvae at 25°C (room temperature (RT) condition) or 4°C (cold temperature (CT) condition). The orange dashed line and solid blue line indicate mean values of absorbance under RT or CT conditions, respectively. Orange and blue shading indicates the 95% confidence intervals of absorbance under RT or CT conditions, respectively. (C) Absorbance at 629 nm of haemolymph collected from *Hayan*^1^ mutant larvae under RT or CT conditions are displayed in box plots. \*\*\**P* < 0.001, Mann-Whitney *U*-test; *n* = 5 in RT and *n* = 5 in CT.

If the contaminated materials were excreted from the larval anus during haemolymph collection, suppression of sphincter muscle contraction by cooling should help reduce such a contamination. To induce and maintain larvae in a state of cold anesthesia, all the equipment and reagents used for measurement including surgical tools, sampling tubes, and distilled water for testing were precooled, and all manipulations were performed at cold temperature (CT, 4.0 ± 2.0°C). We then examined the absorbance at 629 nm and measured the osmolality of the haemolymph collected under CT condition. The optical absorbance at 629 nm was significantly reduced (Figure 2A and 2B; 0.6 ± 0.2, *P* = 3.11 ×10^−4^, Mann-Whitney *U*-test; *n* = 8) and the measured osmolality was 319 ± 5.7 mOsm kg^−1^ (Figure 2B; *P* = 3.11×10^−4^, Mann-Whitney *U*-test; *n* = 8). These results imply that the gut-derived contamination through defecation was significantly inhibited during haemolymph collection under CT conditions; therefore, the osmolality of the haemolymph collected under CT conditions is considered closer to the true value than that of the haemolymph collected at RT. Importantly, insufficient precooling of larvae resulted in varying levels of contamination (data not shown).

We then aimed to estimate the true value of haemolymph osmolality using these data. If the deviations of osmolality largely depend on the amount of contaminating gut-derived materials, the data for the fluid osmolality should be linearly correlated with the corresponding absorbances at 629 nm. In fact, a linear model provided a good fit to these data (Figure 2B; *Y* = 26.1*X* + 306.0, *X* = Absorbance at 629 nm, *Y* = Osmolality; Spearman’s rank correlation: *ρ* = 0.806, *P* = 2.85×10^−4^). Based on this linear relation, we propose that the true value of the haemolymph osmolality is close to 306.0 mOsm kg^−1^, corresponding to the *Y*-intercept of the linear equation. This suggests that quasi-isotonic saline solutions, such as the external saline solution and HL3.1, would be more advantageous options for preserving live specimens in a physiological state, particularly in terms of maintaining the appropriate osmolality (Feng et al., 2004; Li et al., 2023; Xiang et al., 2010). Notably, the proposed value is comparable to the reported haemolymph osmolality in *Drosophila* adult (300∼320 mOsm kg^−1^; Jourjine et al., 2016; Senapati et al., 2019).

Next, we tried to determine the specific step at which gut-derived material contaminated the collected haemolymph. Our suspicion was that the strong centrifugal force during the centrifugation process could have extruded food-derived materials from the hindgut. To test this hypothesis, we tore 10 larvae on a piece of parafilm and harvested the haemolymph leaking from incised sites without centrifugation (Figure 3A), comparing the procedure under RT and CT conditions. The absorbance at 629 nm of haemolymph collected under CT conditions was significantly smaller than with the RT conditions (Figure 3B and 3C; *P* = 0.00794, Man-Whitney *U*-test; *n* = 5), indicating that gut-derived materials had mostly contaminated the sample even before the centrifugation step under RT conditions.

We then asked whether this refined osmolality measurement protocol (CT conditions, with centrifugation, as described in Materials and Methods) is also applicable for a strain that is commonly used as an experimental control (*y*^1^ *w*^67c23^). We measured the osmolality and absorbance at 629 nm of the collected haemolymph under RT and CT conditions and found that the measured values of optical absorbance at 629 nm were significantly different (Figure S2A; [RT] 2.0 ± 0.2, [CT] 1.5 ± 0.2; *P* = 0.0411, Mann-Whitney *U*-test; *n* = 6) and the osmolality values were also significantly different (Figure S2A; [RT] 334 ± 4.3 mOsm kg^−1^, [CT] 310 ± 6.8 mOsm kg^−1^; *P* = 2.17×10^−4^, Mann-Whitney *U*-test; *n* = 6). Here, a linear model provided only a modest fit to the data (Figure S2; *Y* = 24.6*X* + 279.6, *X* = Absorbance at 629 nm, *Y* = Osmolality; Spearman’s rank correlation: *ρ* = 0.587, *P* = 0.049). These findings suggest that our refined measurement protocol is also applicable for other strains with intact melanization ability. Nonetheless, the perturbations in osmolality measurement should be taken into account, and these may be attributed to the significant melanization.

In this study, we refined a method to assess haemolymph osmolality in *Drosophila* larvae. Our new protocol allows us to reliably evaluate the osmolality of small insects, which has previously shown wide variability in reported values, due to the technical difficulty in collecting pure haemolymph from small legless insects. Furthermore, our investigation illuminated the pivotal role of a careful precooling procedure in preventing contamination of the haemolymph by gut-derived materials. In addition, this protocol could be applied for robust measurement of haemolymph osmolality in other small insects such as mosquito larvae or *Leptopilina boulardi* larvae, a parasitoid wasp species that develops inside *Drosophila* larvae.

## Author contributions

Conceptualization, M.K., Y.T., T.Us.; Methodology, M.K., Y.T.; Software, Y.T.; Validation, M.K., Y.T., T.Ue., T.Us.; Formal analysis, M.K., Y.T.; Investigation, M.K., Y.T.; Resources, T.Us.; Data curation, M.K., Y.T.; Writing - original draft, M.K., T.Us.; Writing - review & editing, M.K., Y.T., T.Ue., T.Us.; Visualization, M.K., Y.T.; Supervision, T.Us.; Project administration, T.Us.; Funding acquisition, Y.T., T.Ue., T.Us.

## Data availability

All relevant data can be found within the article and its supplementary information.

## Acknowledgments

We would like to thank M. Futamata, S. Oki, H. Imai, R. Muraki, and Y. Niitani for excellent technical assistance; members of the Uemura lab for experimental advice and discussions; D. Watanabe and S. Yawata for sharing the vapor pressure osmometer; T. Nishimura and N. Okamoto for experimental advice and discussions; J. Hejna for feedback on the manuscript; Won-Jae Lee at Seoul National University for the *Hayan* mutant strain; Bloomington *Drosophila* Stock Center for providing a fly stock. We also thank FlyBase and the *Drosophila* Genomics Resource Center.

---

**Figure S1.**
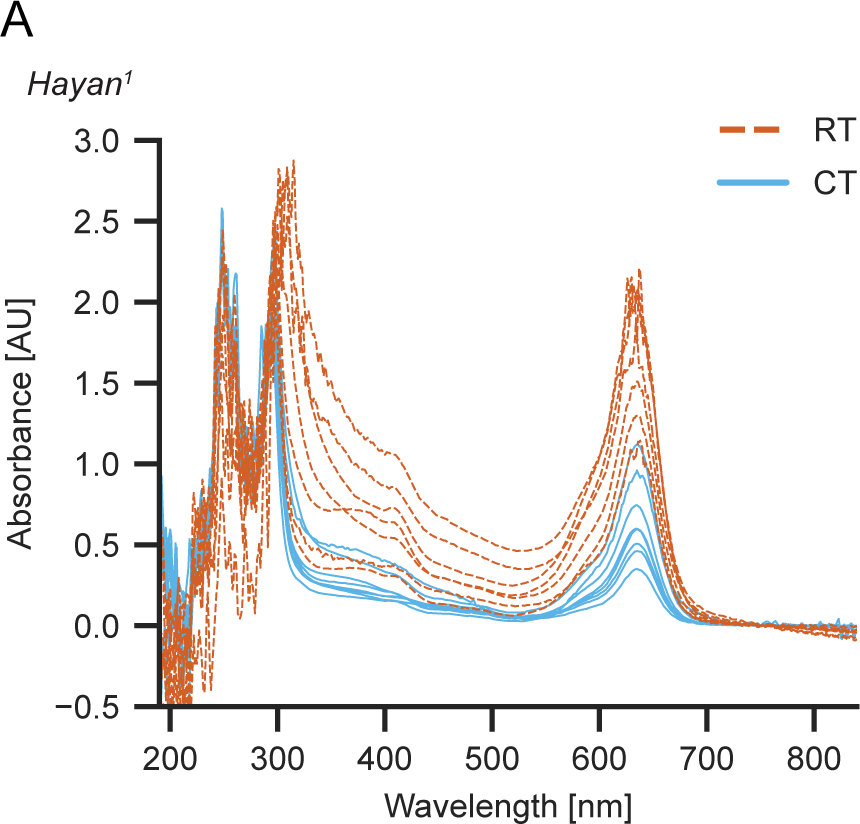
Individual absorption spectra from which the mean values shown in Figure 2A were derived.

**Figure S2.**
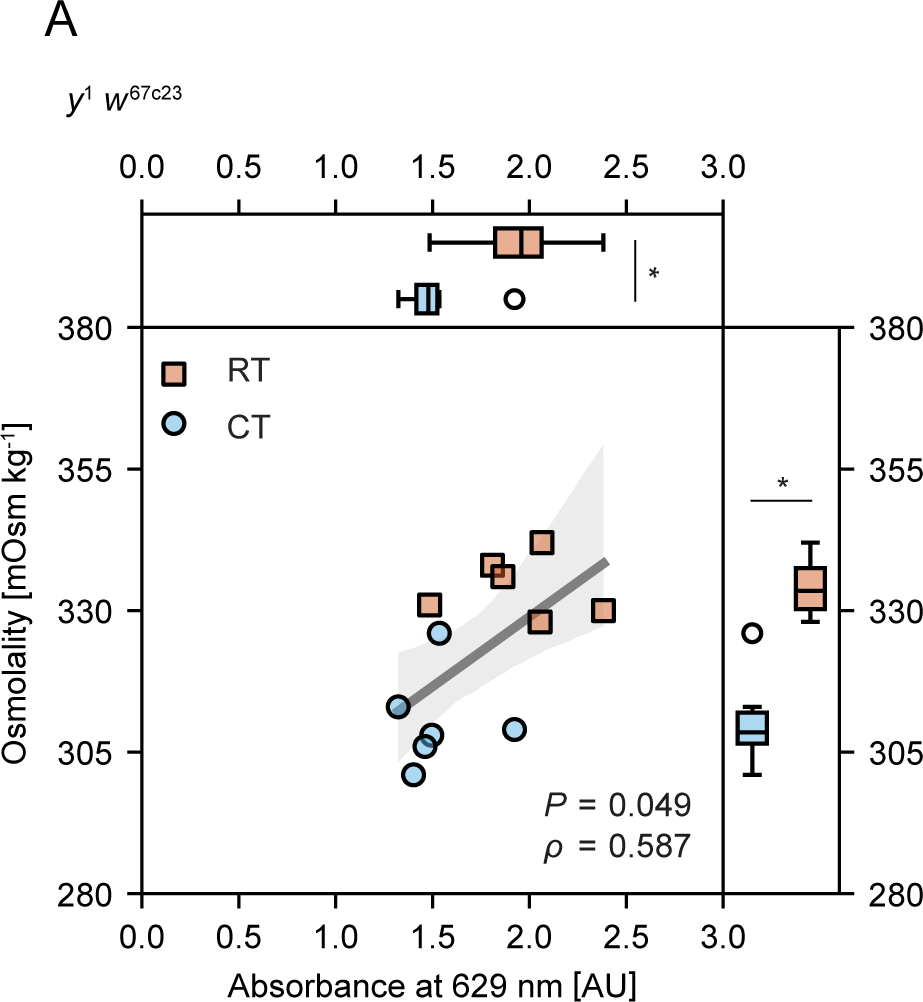
Absorbance at 629 nm and osmolality of haemolymph collected from a commonly used strain. Absorbance at 629 nm and the osmolality of haemolymph collected from *y*^1^ *w*^67c23^ larvae under RT or CT conditions (\**P* < 0.05, \*\**P* < 0.01, \*\*\**P* < 0.001, Mann-Whitney *U*-test; *n* = 6). The dark gray line is a linear regression; light gray shading indicates the 95% confidence interval.

## REFERENCES

Adams, J. R., & Wilcox, T. A. (1973). Determination of Osmolalities of Insect Hemolymph from Several Species. Annals of the Entomological Society of America, 66(3), 575–577. 10.1093/aesa/66.3.575

Borsuk, G., Ptaszyńska, A. A., Olszewski, K., Domaciuk, M., Krutmuang, P., & Paleolog, J. (2017). A New Method for Quick and Easy Hemolymph Collection from Apidae Adults. PLOS ONE, 12(1), e0170487. 10.1371/journal.pone.0170487

Dudzic, J. P., Hanson, M. A., Iatsenko, I., Kondo, S., & Lemaitre, B. (2019). More Than Black or White: Melanization and Toll Share Regulatory Serine Proteases in Drosophila Cell Reports, 27(4), 1050–1061.e3. 10.1016/j.celrep.2019.03.101

Feng, Y., Ueda, A., & Wu, C.-F. (2004). A MODIFIED MINIMAL HEMOLYMPH-LIKE SOLUTION, HL3.1, FOR PHYSIOLOGICAL RECORDINGS AT THE NEUROMUSCULAR JUNCTIONS OF NORMAL AND MUTANT DROSOPHILA LARVAE. Journal of Neurogenetics, 18(2), 377–402. 10.1080/01677060490894522

Handke, B., Szabad, J., Lidsky, P. V., Hafen, E., & Lehner, C. F. (2014). Towards long term cultivation of Drosophila wing imaginal discs in vitro. PLoS ONE, 9(9). 10.1371/journal.pone.0107333

Harpur, R. P., & Popkin, J. S. (1965). OSMOLALITY OF BLOOD AND INTESTINAL CONTENTS IN THE PIG, GUINEA PIG, AND ASCARIS LUMBRICOIDES. Canadian Journal of Biochemistry, 43(7), 1157–1169. 10.1139/o65-128

Hu, Y., Wang, C., Yang, L., Pan, G., Liu, H., Yu, G., & Ye, B. (2020). A Neural Basis for Categorizing Sensory Stimuli to Enhance Decision Accuracy. Current Biology, 30(24), 4896–4909.e6. 10.1016/j.cub.2020.09.045

Jan LY, & Jan YN. (1976). Properties of the larval neuromuscular junction in Drosophila melanogaster. The Journal of Physiology, 262, 189–214.

Jourjine, N., Mullaney, B. C., Mann, K., & Scott, K. (2016). Coupled Sensing of Hunger and Thirst Signals Balances Sugar and Water Consumption. Cell, 166(4), 855–866. 10.1016/j.cell.2016.06.046

Jovanic, T., Schneider-Mizell, C. M., Shao, M., Masson, J.-B., Denisov, G., Fetter, R. D., Mensh, B. D., Truman, J. W., Cardona, A., & Zlatic, M. (2016). Competitive Disinhibition Mediates Behavioral Choice and Sequences in Drosophila. Cell, 167(3), 858–870.e19. 10.1016/j.cell.2016.09.009

Kanaoka, Y., Onodera, K., Watanabe, K., Hayashi, Y., Usui, T., Uemura, T., & Hattori, Y. (2023). Inter-organ Wingless/Ror/Akt signaling regulates nutrient-dependent hyperarborization of somatosensory neurons. ELife, 12, 1–30. 10.7554/eLife.79461

Lemon, W. C., Pulver, S. R., Höckendorf, B., McDole, K., Branson, K., Freeman, J., & Keller, P. J. (2015). Whole-central nervous system functional imaging in larval Drosophila. Nature Communications, 6(May), 7924. 10.1038/ncomms8924

Li, K., Tsukasa, Y., Kurio, M., Maeta, K., Tsumadori, A., Baba, S., Nishimura, R., Murakami, A., Onodera, K., Morimoto, T., Uemura, T., & Usui, T. (2023). Belly roll, a GPI-anchored Ly6 protein, regulates Drosophila melanogaster escape behaviors by modulating the excitability of nociceptive peptidergic interneurons. ELife, 12, 1– 31. 10.7554/eLife.83856

Łoś, A., & Strachecka, A. (2018). Fast and Cost-Effective Biochemical Spectrophotometric Analysis of Solution of Insect “Blood” and Body Surface Elution. Sensors, 18(5), 1494. 10.3390/s18051494

Macleod, G. T., Hegström-Wojtowicz, M., Charlton, M. P., & Atwood, H. L. (2002). Fast calcium signals in Drosophila motor neuron terminals. Journal of Neurophysiology, 88(5), 2659–2663. 10.1152/jn.00515.2002

Nakamizo-Dojo, M., Ishii, K., Yoshino, J., Tsuji, M., & Emoto, K. (2023). Descending GABAergic pathway links brain sugar-sensing to peripheral nociceptive gating in Drosophila. Nature Communications, 14(1), 6515. 10.1038/s41467-023-42202-9

Nam, H.-J., Jang, I.-H., You, H., Lee, K.-A., & Lee, W.-J. (2012). Genetic evidence of a redox-dependent systemic wound response via Hayan Protease-Phenoloxidase system in Drosophila. The EMBO Journal, 31(5), 1253–1265. 10.1038/emboj.2011.476

Onodera, K., Baba, S., Murakami, A., Uemura, T., & Usui, T. (2017). Small conductance Ca2+-activated K+ channels induce the firing pause periods during the activation of Drosophila nociceptive neurons. ELife, 6, 1–17. 10.7554/eLife.29754

Palomino-Schätzlein, M., Carranza-Valencia, J., Guirado, J., Juarez-Carreño, S., & Morante, J. (2022). A toolbox to study metabolic status of Drosophila melanogaster larvae. STAR Protocols, 3(1). 10.1016/j.xpro.2022.101195

Pierce, V. A., Mueller, L. D., & Gibbs, A. G. (1999). Osmoregulation in Drosophila melanogaster selected for urea tolerance. Journal of Experimental Biology, 202(17), 2349–2358. 10.1242/jeb.202.17.2349

Redhai, S., Pilgrim, C., Gaspar, P., Giesen, L. Van, Lopes, T., Riabinina, O., Grenier, T., Milona, A., Chanana, B., Swadling, J. B., Wang, Y.-F., Dahalan, F., Yuan, M., Wilsch-Brauninger, M., Lin, W., Dennison, N., Capriotti, P., Lawniczak, M. K. N., Baines, R. A., … Miguel-Aliaga, I. (2020). An intestinal zinc sensor regulates food intake and developmental growth. *Nature*, May 2019. 10.1038/s41586-020-2111-5

Sano, H., Nakamura, A., Texada, M. J., Truman, J. W., Ishimoto, H., Kamikouchi, A., Nibu, Y., Kume, K., Ida, T., & Kojima, M. (2015). The Nutrient-Responsive Hormone CCHamide-2 Controls Growth by Regulating Insulin-like Peptides in the Brain of Drosophila melanogaster. PLoS Genetics, 11(5), 1–26. 10.1371/journal.pgen.1005209

Schneider, I. (1972). Cell lines derived from late embryonic stages of Drosophila melanogaster. Development, 27(2), 353–365. 10.1242/dev.27.2.353

Senapati, B., Tsao, C.-H., Juan, Y.-A., Chiu, T.-H., Wu, C.-L., Waddell, S., & Lin, S. (2019). A neural mechanism for deprivation state-specific expression of relevant memories in Drosophila. Nature Neuroscience, 22(12), 2029–2039. 10.1038/s41593-019-0515-z

Stewart, B. A., Atwood, H. L., Renger, J. J., Wang, J., & Wu, C.-F. (1994). Improved stability of Drosophila larval neuromuscular preparations in haemolymph-like physiological solutions. Journal of Comparative Physiology A, 175(2), 179–191. 10.1007/BF00215114

Tabunoki, H., Dittmer, N. T., Gorman, M. J., & Kanost, M. R. (2019). Development of a new method for collecting hemolymph and measuring phenoloxidase activity in Tribolium castaneum. BMC Research Notes, 12(1), 7. 10.1186/s13104-018-4041-y

Terada, S.-I., Matsubara, D., Onodera, K., Matsuzaki, M., Uemura, T., & Usui, T. (2016). Neuronal processing of noxious thermal stimuli mediated by dendritic Ca2+ influx in Drosophila somatosensory neurons. ELife, 5, 1–26. 10.7554/eLife.12959

Xiang, Y., Yuan, Q., Vogt, N., Looger, L. L., Jan, L. Y., & Jan, Y. N. (2010). Light-avoidance-mediating photoreceptors tile the Drosophila larval body wall. Nature, 468(7326), 921–926. 10.1038/nature09576

